# Epitranscriptomic addition of m^6^A regulates HIV-1 RNA stability and alternative splicing

**DOI:** 10.1101/2021.02.23.432449

**Authors:** Kevin Tsai, Hal P. Bogerd, Edward M. Kennedy, Ann Emery, Ronald Swanstrom, Bryan R. Cullen

## Abstract

Previous work in several laboratories has demonstrated that the epitranscriptomic addition of m^6^A to viral transcripts promotes the replication and pathogenicity of a wide range of DNA and RNA viruses, yet the underlying mechanisms responsible for this positive effect have remained unclear. It is known that m^6^A function is largely mediated by cellular m^6^A binding proteins or readers, yet how m^6^A readers regulate viral gene expression in general, and HIV-1 gene expression in particular, has been controversial. Here, we confirm that m^6^A addition indeed regulates HIV-1 RNA expression and demonstrate that this effect is in large part mediated by the the nuclear m^6^A reader YTHDC1 and the cytoplasmic m^6^A reader YTHDF2. Both YTHDC1 and YTHDF2 bind to multiple distinct and overlapping sites on the HIV-1 RNA genome, with YTHDC1 recruitment serving to regulate the alternative splicing of HIV-1 RNAs while YTHDF2 binding correlates with increased HIV-1 transcript stability.

**Author Summary:** This manuscript reports that the expression of mRNAs encoded by the pathogenic human retrovirus HIV-1 is regulated by the methylation of a small number of specific adenosine residues. These in turn recruit a nuclear RNA binding protein, called YTHDC1, which modulates the alternative splicing of HIV-1 transcripts, as well as a cytoplasmic RNA binding protein, called YTHDF2, which stabilizes viral mRNAs. The regulation of HIV-1 gene expression by adenosine methylation is therefore critical for the effective and ordered expression of HIV-1 mRNAs and could represent a novel target for antiviral development.

## Introduction

Eukaryotic mRNAs are subject to a range of covalent modifications at the nucleotide level, collectively referred to as epitranscriptomic modifications. The most common epitranscriptomic modification of mammalian mRNAs is methylation of the N^6^ position of adenosine, referred to as m^6^A, which comprises ∼0.4% of all adenosines [1, 2]. m^6^A residues are deposited on mRNAs by a “writer” complex, minimally consisting of METTL3, METTL14 and WTAP, which adds m^6^A to some, but not all, copies of the RNA motif 5’-RRACH-3’ (R=G/A, H=A/U/C) [3–6]. Cells also express two demethylases or “erasers”, called FTO and ALKBH5, which have been proposed to dynamically regulate m^6^A levels [7, 8]. Finally, the phenotypic effects of m^6^A on mRNA metabolism are largely conferred by m^6^A binding proteins, referred to as m^6^A readers, that regulate several steps of RNA metabolism. These include the nuclear m^6^A reader YTHDC1, which has been reported to regulate cellular mRNA splicing and nuclear export, and the cytoplasmic m^6^A readers YTHDF1, YTHDF2 and YTHDF3, which are thought to regulate the translation and stability of m^6^A-containing mRNAs [9–12]. Recent reports suggest that the cytosolic readers YTHDF1, 2 and 3 confer redundant functions, with YTHDF2 the dominant reader due to its higher expression level [13, 14].

While m^6^A is the most common epitranscriptomic modification on cellular mRNAs, analysis of the genomic RNAs encoded by retroviruses revealed an even higher level of m^6^A on these transcripts, with m^6^A comprising as much as ∼2.4% of the adenosines found on murine leukemia virus (MLV) genomic RNAs [15]. This high prevalence suggests that m^6^A might be acting to promote some aspect(s) of the viral replication cycle. Indeed, addition of m^6^A has now been shown to boost the replication of a diverse range of viruses, including not only MLV but also influenza A virus (IAV), SV40, human metapneumovirus (HMPV), enterovirus 71 and respiratory syncytial virus (RSV) [16]. In the case of IAV, HMPV and RSV, the presence of m^6^A residues on viral transcripts was also shown to boost pathogenicity *in vivo*. Similarly, for HIV-1, knock down of the m^6^A writers METTL3 and METTL14 inhibits HIV-1 gene expression and replication while knock down of the m^6^A erasers FTO or ALKBH5 had the opposite effect [17, 18]. However, the mechanism by which this positive effect is achieved has remained controversial. One group [18] reported that a key adenosine residue in the HIV-1 Rev Response Element (RRE) is methylated to m^6^A and that this m^6^A residue promotes Rev binding. However, we and others [17, 19–21] have been unable to confirm the presence of any m^6^A residues on the RRE.

Previously, we reported [20] that the cytoplasmic m^6^A readers YTHDF1, YTHDF3 and especially YTHDF2 all promote HIV-1 gene expression and replication yet another group [17, 22] has argued that these same proteins bind to HIV-1 genomic RNAs and are then packaged into virions where they act to inhibit reverse transcription. In contrast, a third group has recently reported that while YTHDF3 is selectively incorporated into HIV-1 virions, it is then inactivated by cleavage by the viral protease. Only when cleavage by the HIV-1 protease was inhibited was YTHDF3 found to exert an inhibitory effect [23]. Here, we have sought to more fully define how m^6^A residues interact with cellular readers to positively affect HIV-1 mRNA function and report that m^6^A promotes HIV-1 gene expression by at least two distinct mechanisms. On the one hand, m^6^A assures the optimal alternative splicing of HIV-1 transcripts by recruiting the nuclear m^6^A reader YTHDC1. On the other hand, m^6^A also enhances the stability of viral mRNAs by recruiting the cytoplasmic m^6^A reader YTHDF2.

## Results

### Removal of m^6^A residues reduces the level of HIV-1 RNA expression

Previously, it has been demonstrated that knock down of the m^6^A writers METTL3 and METTL14 using RNA interference (RNAi) inhibits HIV-1 Gag expression while knock down of the m^6^A erasers ALKBH5 or FTO increases HIV-1 Gag expression [17, 18], yet the molecular basis for this positive effect of the m^6^A modification has remained unclear. As an alternative approach, we overexpressed the m^6^A eraser ALKBH5, which has been previously demonstrated to globally reduce m^6^A levels on RNA [7, 24]. A lentiviral ALKBH5 expression vector was used to transduce the CD4^+^ T cell line CEM and three single cell clones were isolated, each expressing an ∼3-fold higher level of ALKBH5 than the parental cells (Figs. 1A and 1B). To confirm m^6^A depletion on mRNAs expressed in these ALKBH5 over-expressing cells (+ALKBH5), we quantified the m^6^A content of purified poly(A)^+^ RNA extracted from HIV-1 infected parental or +ALKBH5 CEM cells at 3 days post-infection (dpi) using an m^6^A ELISA assay. Isolation of poly(A)^+^ RNA by oligo-dT purification successfully removed >90% of rRNA (Fig. S1) and we observed that poly(A)+ RNA isolated from the +ALKBH5 CEM cells contained ∼50% less m^6^A than WT CEM cells (Fig. 1C).

**Fig 1.**
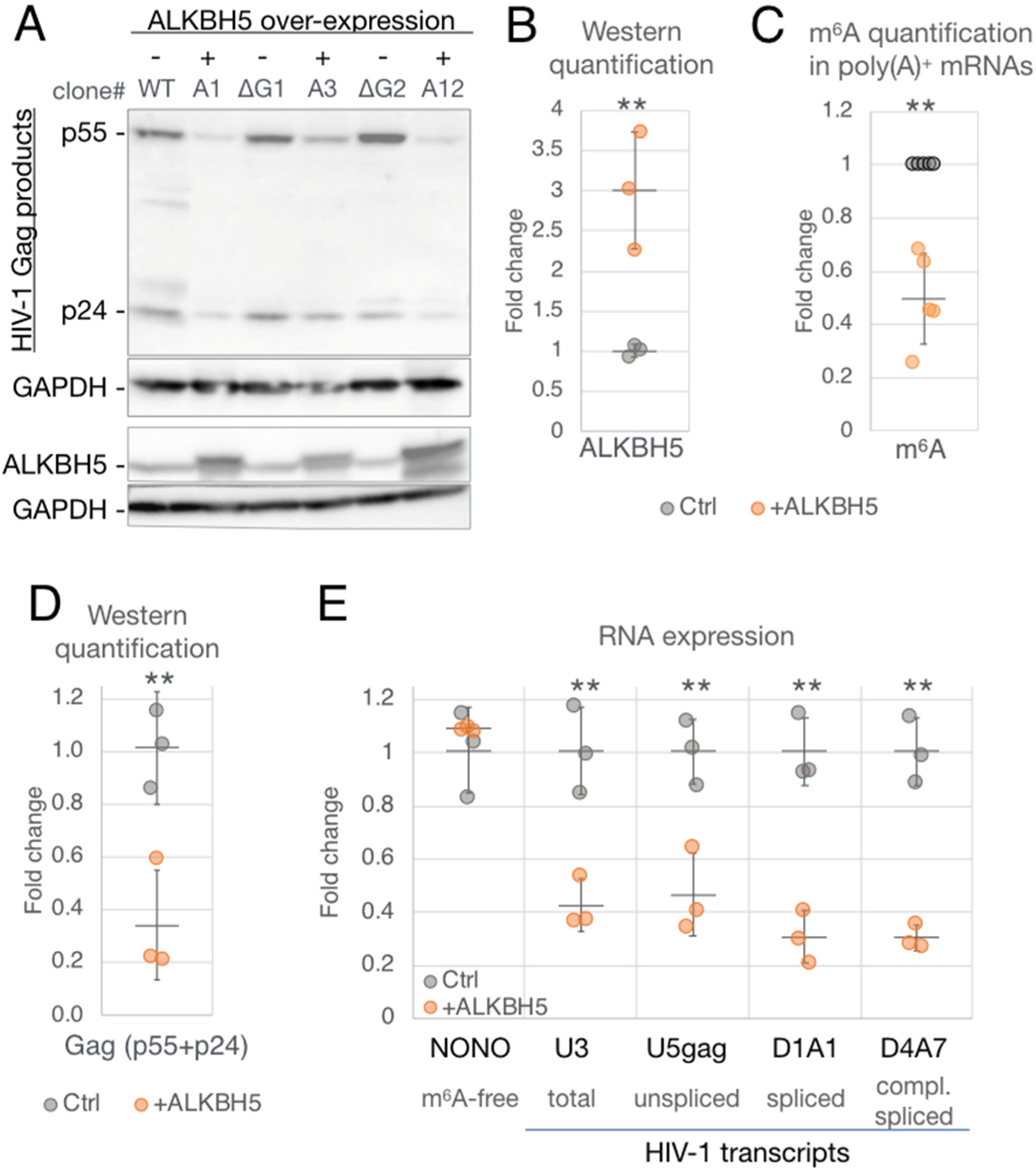
Global depletion of m^6^A in T cells suppresses HIV-1 RNA and protein expression. Three separate clones of the CD4^+^ CEM T cell line (A1, A3 and A12) were obtained by transduction with a lentiviral ALKBH5 expression vector (+ALKBH5). They were then compared to WT CEM cells (Ctrl) for the level of m^6^A present on poly(A)^+^ RNAs. (A) HIV-1 Gag expression levels were analyzed by Western blot at 2 dpi after single cycle HIV-1 infection of +ALKBH5 CEM cells or control cells (one WT and the others expressing the irrelevant GFP-targeted Cas9 (ΔG)). Also analyzed were ALKBH5 (the overexpressed ALKBH5 is epitope tagged and runs slightly slower than endogenous ALKBH5) and GAPDH, as a loading control. (B) ALKBH5 band intensities from panel A were quantified and are given relative to control cells, set at 1. (C) The m^6^A content of poly(A)^+^ mRNAs from Ctrl and +ALKBH5 cells was quantified by ELISA and is given relative to control cells, set at 1 (n=5). (D) Quantification of the HIV-1 Gag band intensities (p24 + p55) from panel A. (E) Aliquotes of the samples from panel C were analyzed for viral RNA expression levels by qRT-PCR, with primers targeting different viral transcripts as marked. The m^6^A-free cellular transcript (NONO) served as a negative control. Statistical analyses by two-tailed Student’s T test, error bars=SD, *p<0.05, **p<0.01.

We next infected WT and +ALKBH5 CEM cells with HIV-1 and then limited this infection to a single cycle by addition of the reverse transcriptase (RT) inhibitor nevarapine at 16 hours post-infection (hpi), followed by harvest at 48 hpi for analysis of viral gene expression. In all three +ALKBH5 clones, reduced m^6^A addition resulted in not only a ∼60% drop in viral Gag protein expression (Figs. 1A and D) but also a similar ∼60% drop in the level of both unsliced and spliced HIV-1 transcripts (Fig. 1E). These data indicate that m^6^A addition to HIV-1 mRNAs results in enhancement of HIV-1 gene expression at the RNA level.

### YTHDF2 binding stabilizes HIV-1 mRNAs

As noted above, while we have previously reported that the cytoplasmic m^6^A readers YTHDF1, YTHDF2 and YTHDF3 increase HIV-1 gene expression and replication, others have argued that they exert an inhibitory effect [17, 20]. As it has recently been reported that all three YTHDF proteins function by similar mechasnisms [13], and because YTHDF2 is the most highly expressed variant in T cells [20], we focused on this cellular protein. We first sought to confirm that YTHDF2 can indeed enhance viral replication by transfecting 293T cells with a plasmid expressing the HIV-1 receptor CD4, along with either an empty plasmid (Ctrl) or a plasmid expressing YTHDF2 (+DF2), and confirmed that this results in YTHDF2 overexpression (Fig. 2A). Control and YTHDF2 overexpressing cells were then infected with a fully replication competent HIV-1 variant expressing nano luciferase (NLuc) in place of Nef. At 24 hpi we observed a significant, ∼3 fold increase in total viral RNA expression in the +DF2 cells when compared to Ctrl cells (Fig. 2B). Analysis of virally-encoded NLuc expression at 24 and 48 hpi revealed an ∼4 fold higher level of NLuc expression at 24 hpi and demonstrated that this indicator virus was, as expected, able to effectively spread through the CD4+ 293T cell culture while maintaining the higher level of NLuc expression seen at 24 hpi.

**Fig 2.**
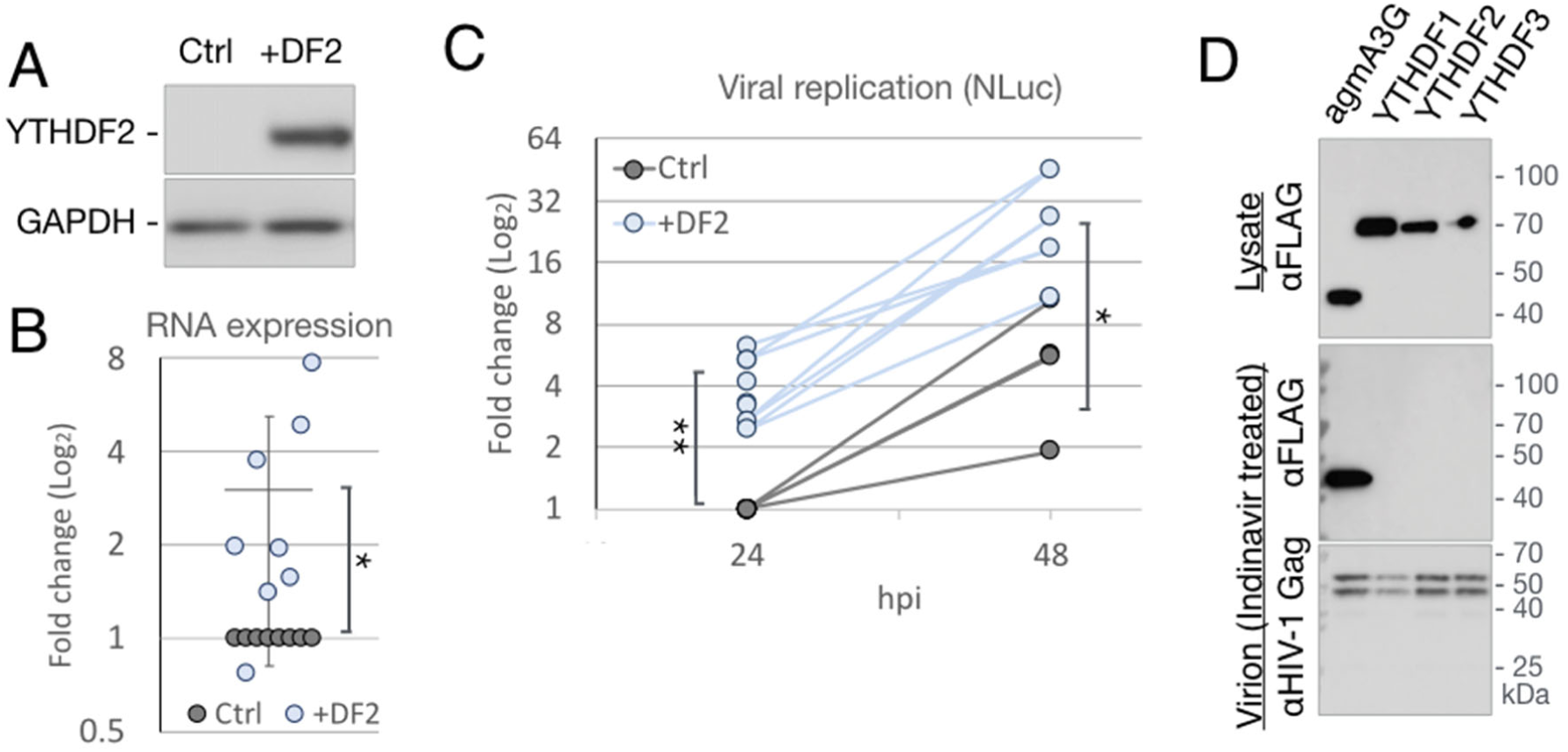
The cytosolic m^6^A reader YTHDF2 enhances HIV-1 replication yet is not packaged into virions. (A-C) 293T cells transfected with either empty vector (Ctrl) or a YTHDF2 expression plasmid (+DF2) were infected with the NL4-3-NLuc reporter virus and collected at 24 hpi for (A) Western blot detection of over-expressed YTHDF2 or (B) qRT-PCR analysis of viral RNA expression, n=8. (C) Quantification of the virally encoded NLuc protein as a measure of viral replication at 24 hpi (n=8) and 48 hpi (n=4). (D) 293T cells were co-transfected with NL4-3-NLuc along with expression vectors expressing FLAG-tagged YTHDF readers or agmA3G and treated with the HIV-1 protease inhibitor indinavir. Lysates of the virus producer cells (upper panel) and purified virions from the supernatant media (two lower panels) were analyzed by Western blot for presence of the FLAG-tagged YTHDF or agmA3G protein and the HIV-1 Gag protein. Statistical analysis by two-tailed Student’s T test, *p<0.05, **p<0.01, error bars=SD.

As noted above, one group has reported that all three cytoplasmic YTHDF readers are incorporated into HIV-1 virions, where they act to inhibit reverse transcription, while a second group has reported that virion packaging is specific for YTHDF3, which is then degraded by the viral protease [17, 22, 23]. To determine whether the YTHDF proteins are indeed incorporated into HIV-1 virions, we co-transfected 293T cells with a molecular clone of HIV-1 along with vectors expressing either FLAG-tagged forms of the m^6^A readers YTHDF1, 2 or 3 or expressing FLAG-tagged African green monkey APOBEC3G (agmA3G), which is known to be packaged into virions [25–27]. While all three YTHDF proteins, along with agmA3G, were detected at comparable levels in virus-producing cells (Fig. 2D, upper panel), only agmA3G was detected in highly purified virion particles isolated from the supernatant media by centrifugation through a sucrose cushion followed by banding on an iodixanol gradient (Fig. 2D, center panel). The failure to detect virion-packaged m^6^A readers could not be explained by their HIV-1 protease-mediated degradation, as the virions were prepared in the presence of the viral protease inhibitor indinavir, as confirmed by the fact that only full length, p55 Gag was detected in these virions (Fig. 2D, lower panel)

We next sought to more precisely define the mechanism by which YTHDF2 enhances HIV-1 gene expression using a more biologically relevant system, ie. single cycle infection by HIV-1 of the CD4^+^ T cell line CEM-SS. YTHDF2 knockout (ΔDF2) or wildtype (WT) CEM-SS cells, and YTHDF2-overexpressing (+DF2) or GFP-expressing (+GFP) CEM-SS cells, have been previously described [20]. WT/ΔDF2 and +GFP/+DF2 CEM-SS cells were infected with HIV-1, RT blocked at 16 hpi using nevarapine to ensure a single cycle infection, and then harvested at 48 hpi. We observed both reduced Gag protein expression in the ΔDF2 cells as well as increased Gag expression in the +DF2 cells, confirming that YTHDF2 indeed enhances viral gene expression in a single-cycle non-spreading infection of T cells (Fig. 3A and 3B). We also observed an ∼80% drop in viral RNA levels in the ΔDF2 cells and an ∼80% increase in viral RNA in the infected +DF2 CEM-SS cells, respectively, when compared to control cells (Fig. 3C), again suggesting that YTHDF2 upregulates viral gene expression at the RNA level. YTHDF2 has been reported to be cytosolic [9], which we confirmed in the CEM-SS T cell line (Fig. S3A) and we therefore hypothesized that YTHDF2 was likely regulating HIV-1 RNA stability rather than synthesis. To test this, we again performed a single cycle infection of the WT and ΔDF2 CEM-SS cells but added the transcription inhibitor Actinomycin D (ActD) at 48 hpi. HIV-1 RNA levels were then quantified by qRT-PCR at multiple time points to monitor the rate of RNA decay. We indeed observed a significantly faster decay of HIV-1 transcripts in the ΔDF2 cells, when compared to WT CEM-SS cells (Fig. 3D), while no difference was observed for a previously reported m^6^A-free host transcript called NONO (Fig. 3E) [1]. These data indicate that YTHDF2 boosts HIV-1 gene expression by enhancing the stability of m^6^A^+^ viral RNA.

**Fig 3.**
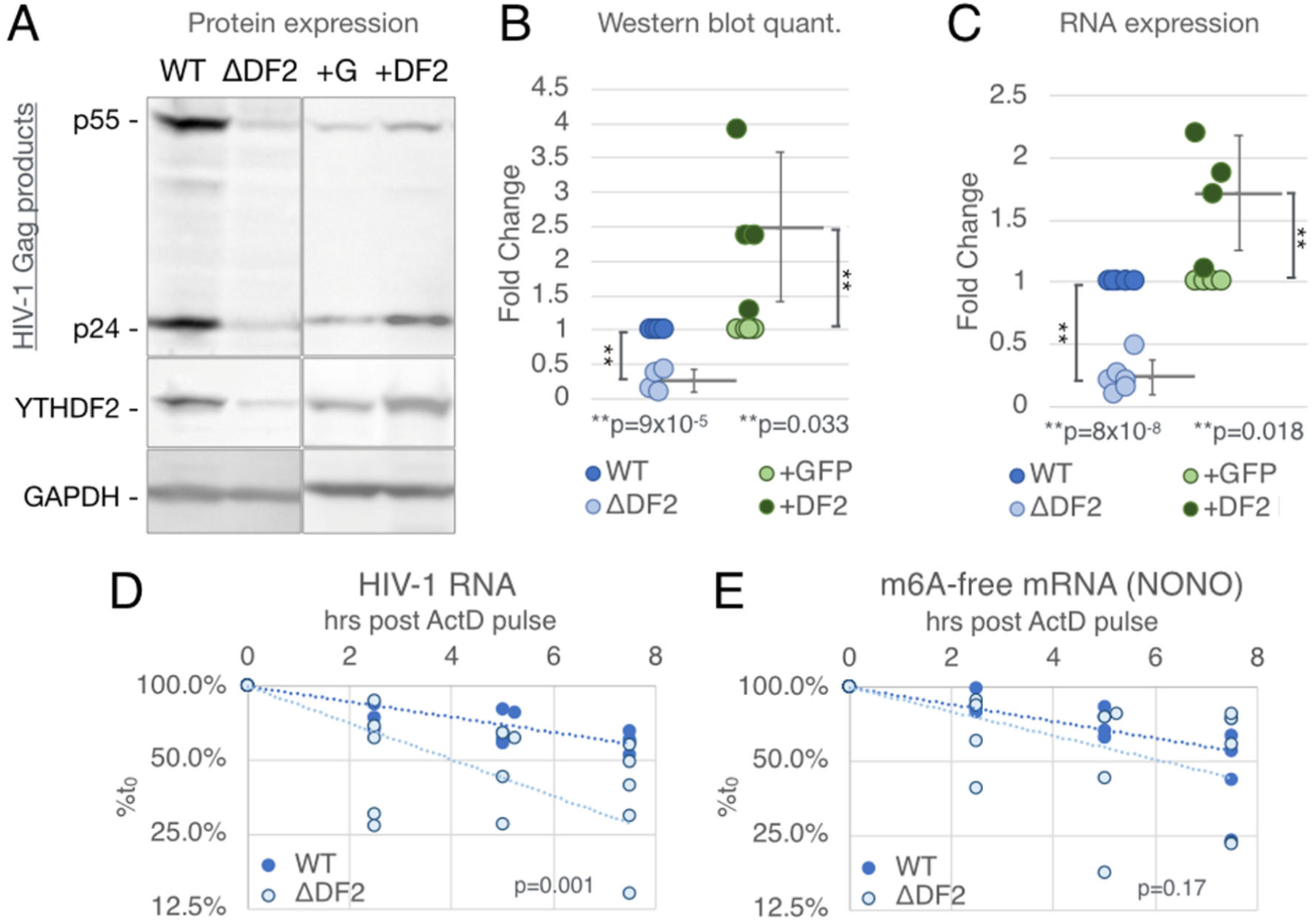
The cytosolic m^6^A reader YTHDF2 enhances HIV-1 RNA stability in CD4^+^ T cells. YTHDF2 knock out CD4^+^ T cells (ΔDF2), WT control cells (WT), and cells transduced with a lentiviral YTHDF2 or GFP expression vector (+DF2 or +G) were used for the following single cycle infection assays. (A) Viral Gag protein expression levels analyzed by Western blot. (B) Quantification of protein band intensities of Western blots as shown in panel A (n=4) (C) Viral RNA expression assayed by qRT-PCR (WT & ΔYTHDF2 n=6, +GFP & +YTHDF2 n=5). (D-E) Viral RNA stability assayed by treating infected WT or ΔDF2 cells at 2 dpi with ActD and collecting RNA at the indicated time points. (D) Viral RNA levels analyzed by qRT-PCR and shown as % of the RNA level at time point 0. (E) RNA stability of a m^6^A-free host transcript (NONO) assayed as a control. Statistical analysis of data in B and C used the two-tailed Student’s T test, p values as marked, error bars=SD. For panels F and G, slopes of regression lines were compared by ANCOVA, n=5.

### Both YTHDC1 and YTHDF2 bind to m^6^A sites on HIV-1 transcripts

HIV-1 replication is tightly regulated by a remarkably complex pattern of alternative splicing that results in the ordered expression of up to 30 unspliced, partially spliced and fully spliced viral mRNAs [28, 29]. Both m^6^A and the nuclear m^6^A reader YTHDC1 have been reported to affect cellular mRNA splicing [1, 10] and we therefore sought to determine if HIV-1 RNA splicing is also epitranscriptomically regulated by YTHDC1 binding to viral m^6^A residues. Initially, we used the photo-assisted cross linking and immunoprecipitation (PAR-CLIP) technique [30] to map YTHDC1 binding sites on HIV-1 transcripts in infected cells and compared these to the previously mapped binding sites for YTHDF1 and YTHDF2 as well as sites of m^6^A addition mapped by antibody binding (PA-m^6^A–seq, Fig. 4) [19, 20]. We identified at least 7 reproducible YTHDC1 binding sites across two independent PAR-CLIP experiments, and found that these largely coincided with previously mapped sites of m^6^A addition [19] as well as with known YTHDF1 and YTHDF2 binding sites [20]. Moreover, these sites all contained the m^6^A addition motif 5’-RRACH-3’ [1, 2]. Due to splicing, the read depth in the first ∼7500nt of the viral genome is markedly lower than in the 3’ untranslated region of viral mRNAs in infected cells. Nevertheless, we were able to map a YTHDC1 binding site adjacent to HIV-1 splice acceptor A3 (Fig. 4A), as well as a binding site overlapping splice acceptor A7 (Fig. 4B), in addition to YTHDC1 binding sites coincident with the previously reported m^6^A sites in the *env/rev* overlap, the *nef* gene and in the TAR hairpin [20]. Conversely, the m^6^A sites at ∼8,900 nt located in the NF-kB sites in the viral LTR U3 region are at best weakly bound by YTHDC1 (Fig. 4B). Of note, we observed no binding of YTHDC1, YTHDF1, YTHDF2 or the m^6^A-specific antibody to the reported m^6^A addition site in the RRE [18], which agrees with m^6^A mapping data on HIV-1 RNAs reported by others [17].

**Fig 4.**
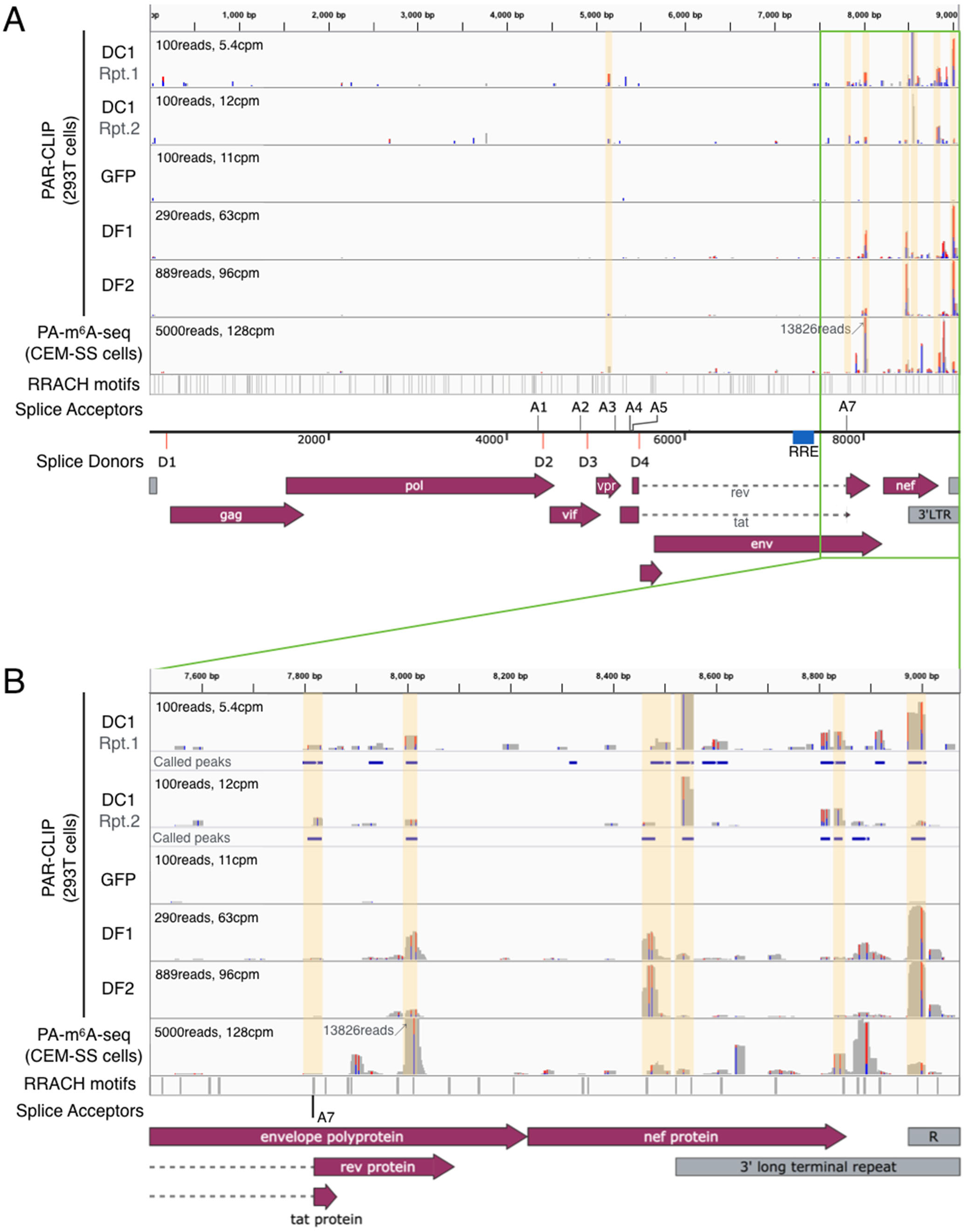
The nuclaer m^6^A reader YTHDC1 binds HIV-1 RNA at previously mapped m^6^A sites. PAR-CLIP was performed on 293T cells transfected with FLAG-GFP or FLAG-YTHDC1, and infected with HIV-1, by immunoprecipitating protein-RNA complexes using a FLAG antibody. Sequencing reads were mapped to the HIV-1 genome, with two independent repeats of YTHDC1 PAR-CLIP (DC1 lanes) shown alongside previously published YTHDF1 & YTHDF2 PAR-CLIP (DF1 & DF2 lanes) and m^6^A mapping results (PA-m^6^A-seq lane) [19, 20]. An overview of the whole HIV-1 genome is shown in (A), with the 3’end (3’ of 7,500bp) shown in (B) where the major m^6^A sites are located. Locations of the m^6^A motif 5’**-**RRACH-3’ are shown in the bottom lane, with a diagram of the splice donors and acceptors relative to the HIV-1 genome map shown beneath. Significant YTHDC1 peaks called by PARalyzer shown as blue bars beneath each DC1 lane in (B). PARalyzer-called peaks that overlap with m^6^A sites and 5’-RRACH-3’ motifs are highlighted in yellow.

### YTHDC1 regulates the expression of HIV-1 transcripts

Having confirmed that YTHDC1 indeed binds m^6^A residues on HIV-1 transcripts in infected cells, we next asked whether YTHDC1 can regulate HIV-1 RNA expression. Because we were unable to obtain YTHDC1 knockout cells, implying that YTHDC1 may be an essential gene, we instead used RNAi to knockdown YTHDC1 expression. We transfected 293T cells with a plasmid encoding the HIV-1 receptor CD4 and also transfected the cells on two successive days with siRNAs targeting YTHDC1 mRNA (siDC1) or with control siRNAs (siCtrl). As expected, we observed a decrease in YTHDC1 protein expression only in the siDC1 transfected cells and this correlated with a surprising increase in HIV-1 Gag protein expression (Fig. 5A, lanes 1-3). If the increase in viral Gag expression is indeed a direct consequence of reduced YTHDC1 expression, then one might predict a decrease in viral Gag expression upon overexpression of YTHDC1 (+DC1), and this was indeed observed (Fig. 5A, compare lanes 4 and 5). A similar effect was seen at the RNA level, with quantification of total, unspliced and spliced viral RNA (specifically viral RNAs spliced from D1 to A1) using qRT-PCR revealing that knock down of YTHDC1 using RNAi resulted in a significantly higher level of viral RNA expression, whereas YTHDC1 overexpression resulted in reduced viral RNA expression (Fig. 5B). This effect was not observed for the NONO transcript lacking m^6^A sites and was more prominent for the spliced viral RNA bearing the D1/A1 splice junction than for the unspliced viral RNA, thus suggesting that the nuclear YTHDC1 protein (Fig. S2B) might regulate HIV-1 RNA expression post-transcriptionally.

**Fig 5.**
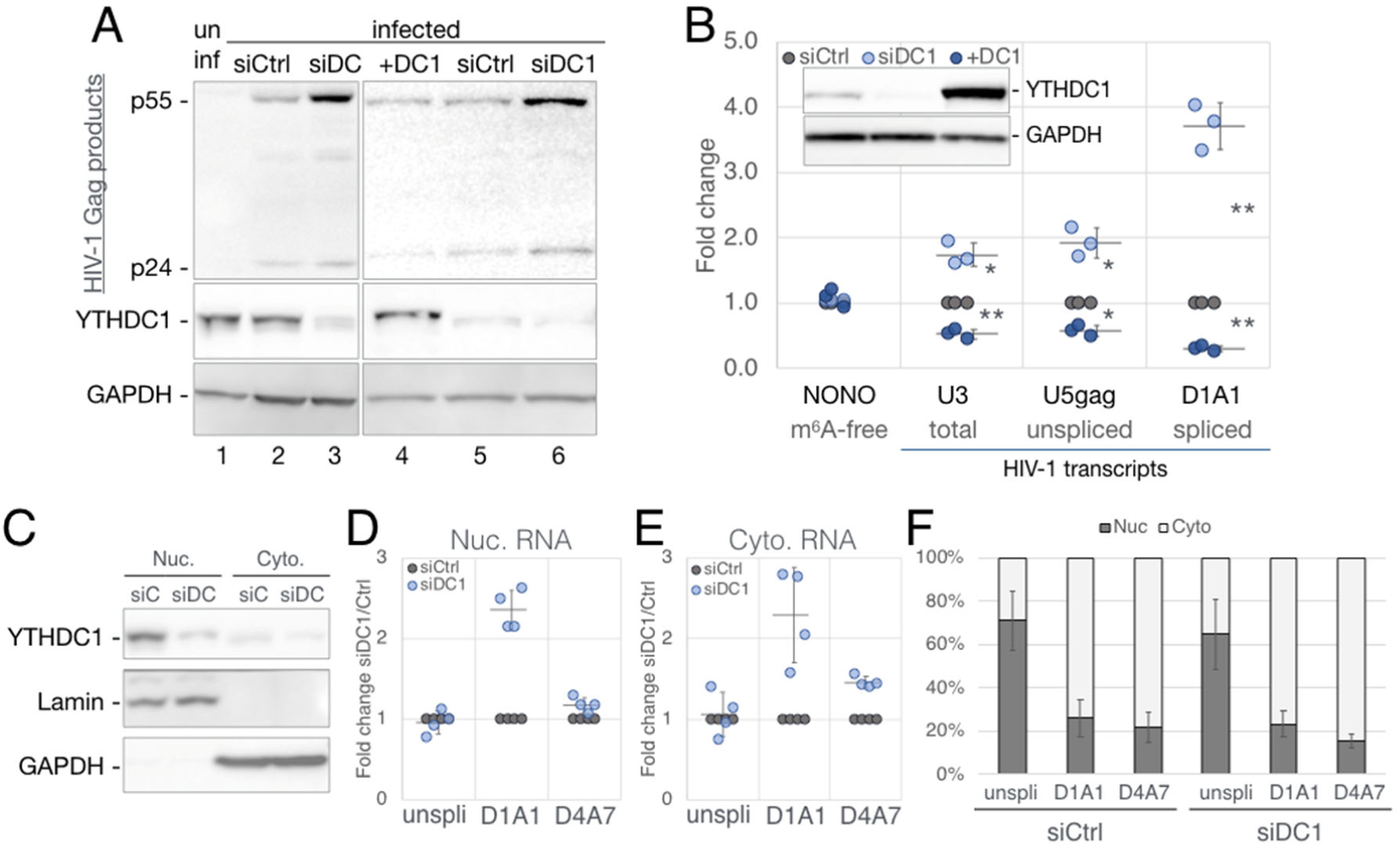
YTHDC1 regulates HIV-1 gene expression with no detectable effect on RNA nuclear export. 293T cells transfected with non-targeting (siCtrl, gray) or YTHDC1-targeting (siDC1, light blue) siRNAs were co-transfected with a YTHDC1 expression vector (+DC1, dark blue) or empty vector. HIV-1 single cycle infections were performed for the following analyses: (A) Viral Gag protein expression assayed by Western blot, co-stained with a YTHDC1 antibody. (B) Viral RNA levels assayed by qRT-PCR, n=3, with the m^6^A-free host NONO mRNA as a control. A representive Western blot shown in the top left inset depicts the validation of YTHDC1 levels for the samples used in this panel. (C-F) Subcellular fractionation assay of infected siCtrl and siDC1 cells. (C) Western blot validation of fractionation, stained for YTHDC1, nuclear Lamin A/C, and the cytosolic protein GAPDH. (D-F) HIV-1 transcript alternative splice forms were quantified by qRT-PCR with primers targeting unspliced (unspli, U5-gag), and the D1/A1, and D4/A7 splice junctions, calculated as fold change of siDC1 over siCtrl in the nuclear (D), and cytosolic fraction (E). (F) The same RNA quantification as panels (D-E) calculated as percent nuclear and percent cytoplasmic. Statistical analysis used Student’s T test, error bars=SD, *<0.05, **<0.01.

### YTHDC1 does not regulate the nuclear export of HIV-1 transcripts

YTHDC1 has been proposed to regulate both the alternative splicing and nuclear export of cellular m^6^A methylated mRNAs [10, 12] and we first asked whether YTHDC1 might regulate the nuclear export of HIV-1 transcripts. To determine whether knock down of YTHDC1 expression had any effect on the nucleocytoplasmic localization of either fully spliced or incompletely spliced HIV-1 RNAs, we isolated nuclear or cytoplasmic fractions from WT or YTHDC1 knock down 293T cells. Analysis of these fractions by Western analysis for the nuclear protein Lamin or the cytoplasmic protein GAPDH confirmed the integrity of the fractionation process and further confirmed not only the nuclear localization of YTHDC1 but also the effective knock down of YTHDC1 protein expression by RNAi (Fig. 5C). Analysis of viral RNA expression in the isolated nuclear and cytoplasmic fractions confirmed that knock down of YTHDC1 expression substantially enhanced the expression of HIV-1 transcripts bearing the D1/A1 splice junction (Figs. 5D and E), as previously noted in total cellular RNA (Fig. 5B). However, this analysis did not reveal any changes in the subcellular localization of either unspliced viral RNA, or fully spliced viral RNAs bearing the D4/A7 splice junction, when YTHDC1 expression was inhibited (Fig. 5D-F).

### YTHDC1 regulates the alternative splicing of HIV-1 transcripts

The observation that knock down of YTHDC1 expression using RNAi strongly enhanced the expression of HIV-1 transcripts bearing the D1/A1 splice junction, while only modestly affecting both unspliced and total viral RNA expression (Fig. 5) suggested that YTHDC1 might regulate the alternative splicing of HIV-1 transcripts, as has indeed been previously proposed for cellular RNAs [10]. To determine whether this is the case, we used Primer-ID-based deep sequencing of splice forms [31] to quantitatively measure the effect of YTHDC1 on HIV-1 alternative splicing. In this assay, a common forward primer that anneals 5’ to the *gag* gene can be paired with either random reverse primers, or splice form-specific 4kb or 1.8kb reverse primers, to selectively amplify viral transcripts (Fig. 6A). HIV-1 transcripts are divided by size into the ∼9kb unspliced transcript, ∼4kb incompletely spliced transcripts (which retain the D4/A7 *env* intron), and ∼1.8kb fully spliced transcripts, (lacking the D4/A7 intron). While the random reverse primer amplifies all viral transcripts, the 4kb reverse primer, located within the D4/A7 intron, only amplifies incompletely spliced transcripts. The 1.8kb reverse primer spans the D4/A7 splice junction and thus only amplifies fully spliced transcripts. Using the random reverse primer, we first noted that ∼33% of viral transcripts are spliced in the siCtrl cells, while this increased to ∼42% in the siDC1 cells but decreased to ∼25% in the +DC1 cells (Fig. 6B). Yet, among spliced transcripts, a constant ∼85% of all spliced transcripts were fully spliced, that is lacking the D4/A7 intron, regardless of YTHDC1 expression level (Fig. 6C). Thus, YTHDC1 has a stronger inhibitory effect on utilization of splice donor D1 than D4. Quantification of the level of splicing between donor D1 and the central splice acceptors A1-A5 (Fig. 6D), using the random reverse primer, revealed that YTHDC1 overexpression led to a significant bias towards utilization of the more 3’ acceptors A3, A4 and A5, at the expense of the more 5’ acceptors A1 and A2 (Fig. 6D) and, as expected, the opposite result was observed in cells in which YTHDC1 expression was knocked down, ie. increased utilization of A1 and A2 and a concomitant reduction in the utilization of A3, A4 and A5. The class-specific 1.8 and 4kb reverse primers confirmed this bias (Figs. 6E and F). YTHDC1 over-expression also led to a significant, ∼2.5x increase in the detection of the normally very rare D1 to A7 splice (Fig. 6G), consistent with the observed bias towards splicing D1 to more 3’ splice acceptors in YTHDC1 overexpressing cells. Finally, we noted that among the transcripts that used splice acceptor A2, YTHDC1 significantly decreased the proportion of viral RNAs that were additionally spliced using splice donor D3 (Fig. 6H), thus incorporating the small non-coding exon located between A2 and D3 (Fig. 6A).

**Fig 6.**
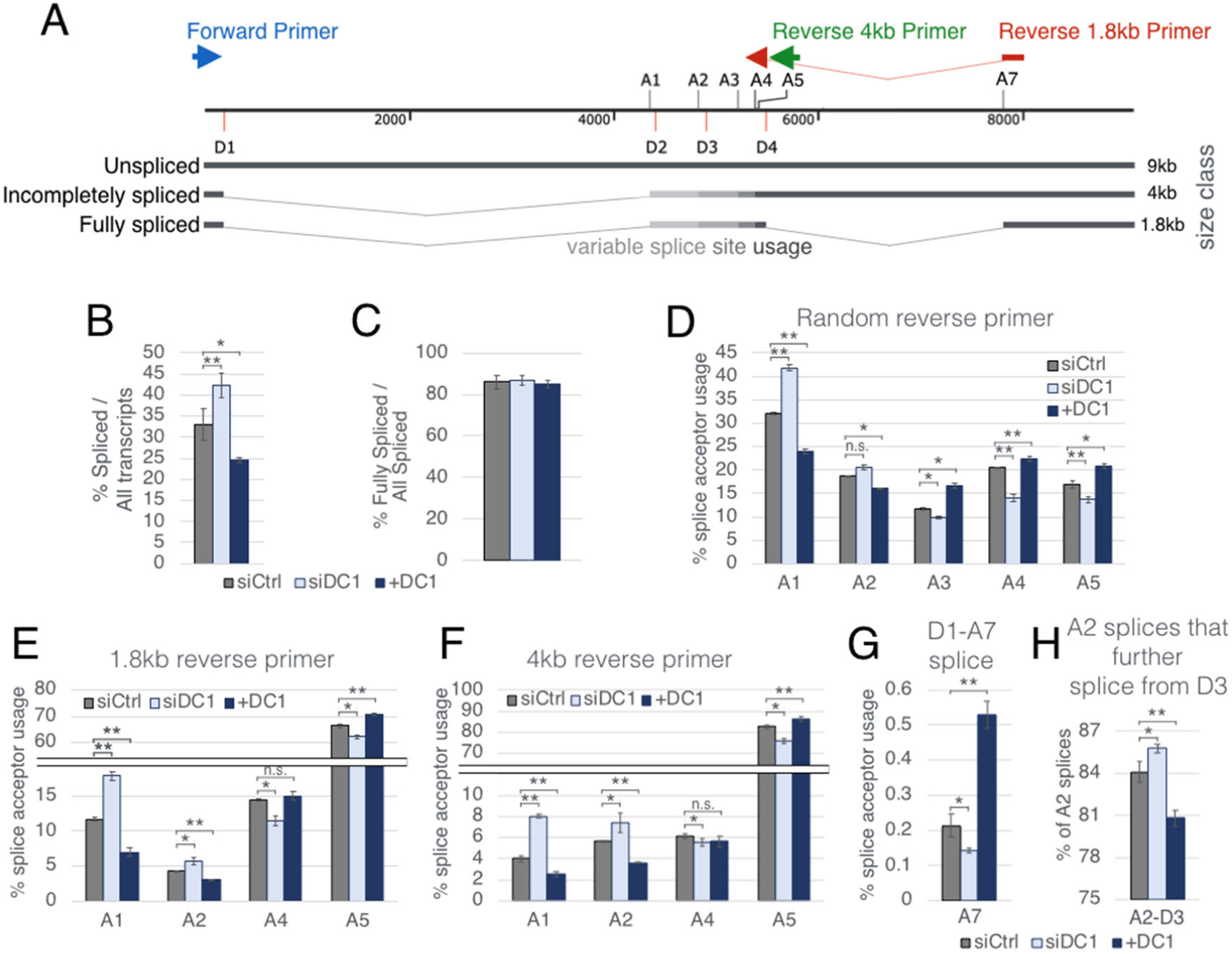
YTHDC1 regulates the relative expression of alternatively spliced HIV-1 RNAs. Viral transcript spliced isoforms in infected siCtrl, siDC1, and +DC1 cells 48 hpi were analyzed by PrimerID RNA-seq, n=3 [31]. (A) Schematic of HIV-1 splice donors and acceptors relative to the three major classes of spliced viral RNAs. Top arrows depict the locations of primers used to amplify sequences for RNA-seq, including the common 5’ forward primer in blue, the reverse 4kb primer in green, and the reverse 1.8kb primer in red spanning the D4/A7 splice junction. A random reverse primer was also used but is not shown. (B) Spliced transcripts are given as a percentage of all transcripts (1.8kb+4kb class read counts / total read counts). (C) Percent of fully spliced transcripts over all spliced transcripts (1.8kb / (1.8kb+4kb) read counts). (D-F) Splice acceptor usage assayed using the common forward primer in conjunction with the random reverse primer (D), 1.8kb reverse primer (E) and 4kb reverse primer (F). Use of acceptor A3 results in long transcripts that are biased against when amplifying using the 1.8 or 4kb reverse primers, A3 usage is thus only shown with the random reverse primer. (G) Percent occurance of D1/A7 splices. (H) Percent occurance of A2-utilizing spliced RNAs that subsequently splice from D3. Statistical analysis used Student’s T test, error bars=SD, *<0.05, **<0.01.

## Discussion

Previously, knock down of the m^6^A writers METTL3 and METTL14 using RNAi was reported to inhibit HIV-1 gene expression, while knock down of the m^6^A erasers FTO and ALKBH5 enhanced HIV-1 gene expression [17, 18]. The resultant conclusion that m^6^A must facilitate one or more aspects of the HIV-1 replication cycle is consistent with the finding that HIV-1 genomic RNAs contain higher levels of m^6^A than the average cellular mRNA [19]. More recently, m^6^A was shown to enhance the replication and pathogenicity of a wide range of DNA and RNA viruses [16], thus implying that the underlying mechanism of action of m^6^A, though still unclear, positively affects the expression and/or function of a range of viral RNA transcripts.

In one early study looking at how m^6^A promotes HIV-1 gene expression, a key adenosine residue on the RRE was proposed to be methylated to m^6^A and then promote Rev binding and HIV-1 RNA nuclear export [18]. However, several later studies failed to detect any m^6^A residues on the RRE [17, 19, 20] and the presence or absence of m^6^A at this RRE residue was found to not, in fact, affect Rev binding [21]. Moreover, as the expression of mRNAs encoded by several cytoplasmic RNA viruses is also boosted by m^6^A addition [16], nuclear RNA export cannot be the primary mechanism of action of m^6^A.

While m^6^A has the potential to affect RNA secondary structure, it is nevertheless clear that the phenotypic consequences of m^6^A addition are largely mediated by m^6^A readers, which include the nuclear m^6^A reader YTHDC1 and the cytoplasmic m^6^A readers YTHDF1, YTHDF2 and YTHDF3. While we have previously reported that all of the cytoplasmic readers, including especially YTHDF2, significantly boost HIV-1 RNA expression in infected cells [20], others have proposed that the YTHDF proteins are bound to m^6^A residues on the HIV-1 genomic RNA and then packaged into HIV-1 virions where they act to inhibit reverse transcription [17]. This latter result is clearly inconsistent with the finding, by this group and others [17, 18, 20], that m^6^A strongly promotes HIV-1 replication. Moreover, as m^6^A is added to RNAs at the consensus sequence 5’-RRACH-3’, mutants lacking this motif should be rapidly selected during passage of a rapidly replicating, error prone RNA virus such as HIV-1 if m^6^A indeed exerted an inhibitory effect in *cis*. Recently it was been reported that, if YTHDF proteins are indeed packaged into HIV-1 virions, then they are effectively inactivated due to degradation by the viral protease [23]. However, we were unable to detect the virion packaging of any YTHDF reader protein even when viral protease activity was inhibited (Fig. 2D).

In this report, we have sought to define the precise step(s) in the HIV-1 replication cycle that is regulated by m^6^A-bound reader proteins. In the case of YTHDF2, we show that this reader primarily acts to increase HIV-1 RNA expression, and not translation, and demonstrate that this effect is mediated by enhanced viral RNA stability (Fig. 2 and 3) This is a surprising result given that most previous studies looking at YTHDF2 function have reported that YTHDF2 binding to cellular mRNAs actually destabilizes the RNA [9, 32], although several reports have suggested that m^6^A can stabilize cellular transcripts in some cases [33] with one report demonstrating that m^6^A addition stabilizes specific cellular mRNAs during hypoxia [34]. We also note that m^6^A is known to promote the expression of mRNAs encoded by a wide range of viruses [16], which is clearly inconsistent with the idea that the recruitment of cytoplasmic readers inhibits mRNA expression. Why YTHDF2, and potentially the other cytoplasmic m^6^A readers, stabilize m^6^A-containing viral RNAs yet destabilize most m^6^A-containing cellular RNAs is currently unknown, though it appears possible that this is dependent on the simultaneous recruitment of other RNA binding proteins that remain to be identified, perhaps to adjacent sites on the viral RNA. We note that the inhibitory effect on HIV-1 gene expression in infected T cells seen when expression of the YTHDF2 reader is blocked (Fig. 3) is indistinguishable from the inhibitory effect seen when removal of m^6^A residues from viral transcripts is enhanced by overexpression of the m^6^A eraser ALKBH5 (Fig. 1), thus arguing that the boost in total viral RNA expression that results from the addition of m^6^A residues is largely mediated by cytoplasmic RNA readers.

While there has been little previous work looking at the effect of m^6^A on viral RNA splicing, with the exception of a recent paper showing that depletion of the m^6^A writer METTL3 reduces the efficiency of splicing of adenoviral late mRNAs [35], it is known that m^6^A addition can regulate cellular mRNA splicing, acting predominantly via the nuclear m^6^A reader YTHDC1 [1, 10, 36, 37]. As HIV-1 transcripts are among the most extensively alternatively spliced RNAs ever identified [29, 38], we were interested in whether YTHDC1 regulated this process in infected cells. Using the PAR-CLIP technique, we were indeed able to map at least 7 distinct YTHDC1 binding sites on HIV-1 transcripts that largely, but not entirely, coincided with previously mapped YTHDF1 and YTHDF2 binding sites as well as sites of m^6^A addition previously mapped using the antibody-dependent technique PA-m^6^A-seq [19, 20] (Fig. 4). As we were unable to knock out YTHDC1 using CRISPR/Cas, we decided to knock down YTHDC1 expression using RNAi, and also overexpress YTHDC1 by transfection, to determine whether changing the level of YTHDC1 expression affected HIV-1 RNA expression and/or alternative splicing. As shown in Figs. 5A and B, we unexpectedly observed that knock down of YTHDC1 actually increased HIV-1 Gag protein and RNA expression while overexpression of YTHDC1 inhibited HIV-1 gene expression. This effect, which is the exact opposite of what we saw when YTHDF2 was knocked out or overexpressed (Figs 2 and 3), was particularly significant for viral RNAs spliced from donor D1 to acceptor A1 (Fig. 5B), thus suggesting that YTHDC1 indeed regulated some aspect(s) of HIV-1 RNA metabolism. Despite previous reports indicating that YTHDC1 can regulate the nuclear export of cellular mRNAs [12], and our report that m^6^A regulates the nuclear export of SV40 transcripts [39], we did not detect any effect of YTHDC1 knock down on the nuclear export of either unspliced HIV-1 transcripts, which is dependent on the viral Rev protein and its cellular cofactor CRM1, or on the subcellular localization of fully spliced viral transcripts bearing the D4/A7 splice junction, which is mediated by the canonical cellular nuclear mRNA export factor NXF1 (Fig. 5) [40]. However, using the previously described “Primer-ID-based deep sequencing of splice forms” technique [31], we were able to document clear and opposite effects of YTHDC1 knock down or overexpression on the relative utilization of specific HIV-1 splice acceptors. Specifically, knock down of YTHDC1 modestly increased viral RNA splicing (Fig. 6B) by increasing the utilization of splice acceptors A2 and particularly A1, while simultaneously reducing the utilization of splice acceptors A3, A4 and A5 (Figs. 6D-F), thus explaining the increased level of D1 to A1 spliced RNA detected by qRT-PCR in Fig. 5. Overexpression of YTHDC1, in contrast, reduced utilization of A1 and A2 while simultaneously increasing the utilization of acceptors A3, A4 and A5, an effect that was most clearly seen when the random reverse primer splicing assay was used (Fig. 6D).

Previous work looking at the effect of YTHDC1 on splicing of cellular mRNAs largely looked at the effect of YTHDC1 on the inclusion or skipping of cassette exons and concluded that YTHDC1 promoted exon inclusion by recruiting the pre-mRNA splicing factor SRSF3 [10]. However, this report did not address whether the location of m^6^A addition sites relative to splice sites was important. As alternative splicing of HIV-1 RNAs is a much more complex process than simple exon inclusion or exclusion, it is hard to address whether our data are consistent with this earlier report. However, we note that none of the mapped YTHDC1 binding sites on the HIV-1 genome colocalize with either known viral splicing enhancer/repressor elements or with any intronic branch points (consensus motif 5’-YNYURAY-3’) [41]. A minor peak of YTHDC1 binding does precisely overlap with splice acceptor A7, with the only m^6^A motif present in this peak (5’-RRm^6^ACH-3’) positioning the splice junction exactly 5’ to the m^6^A (Fig. 4B). YTHDC1 overexpression also biases D1-originating splices away from acceptors A1 and A2, and towards A3, A4, A5 and A7 (Fig. 6) and we mapped a YTHDC1 binding site between A2 and A3 (Fig. 4A), which demarcates the border of this effect. However, determining whether the precise location of the YTHDC1 binding sites mapped on the HIV-1 genome, relative to splice sites, is indeed important may be challenging, given the difficulty of generating silent mutations at many of these sites.

## Materials and methods

### Cell lines and virus clones

HEK293T cells (referred to as 293T), a human kidney epithelial cell line of female origin, was purchased from the American Type Culture Collection. CEM and CEM-SS cells are human CD4^+^ T cell lines of female origin, with CEM-SS a subclone of CEM. Both were obtained from the NIH AIDS reagent program. 293T cells were cultured in Dulbecco’s Modified Eagle’s Medium (DMEM) with 6% fetal bovine serum (FBS) and 1% Antibiotic-Antimycotic (Gibco, 15240062). T cell lines were cultured in Roswell Park Memorial Institute (RPMI) 1640 medium supplemented with 10% FBS and 1% Antibiotic-Antimycotic.

The +ALKBH5 cells were produced by transducing CEM cells with a lentiviral vector expressing FLAG-ALKBH5 followed by puromycin selection and single cell cloning. YTHDF2 knock out and overexpressing CEM-SS cells were previously described [20]. YTHDC1 mRNA knockdown in 293T cells was performed using siRNAs (Origene # SR314128) transfected using Lipofectamine RNAiMax (Invitrogen).

Recombinant virus clones used include the laboratory strain NL4-3, obtained from the NIH AIDS Reagent [42], and the nano-luciferase reporter virus NL4-3-NLuc, in which the viral *nef* gene has been substituted with the NLuc indicator gene [43].

### Antibodies

Anti-FLAG (M2), Sigma F1804. Anti-HIV-1 p24 gag #24-3, ARP-6458. Anti-GAPDH, Proteintech 60004-1-Ig. Anti-ALKBH5, Proteintech 16837-1-AP. Anti-YTHDF2, Proteintech 24744-1-AP. Anti-YTHDC1, Abcam ab122340. Anti-mouse IgG, Sigma A9044. Anti-rabbit IgG, Sigma A6154.

### Cloning of protein expression plasmids

ALKBH5 was cloned out of cDNA produced from CEM cells, by PCR with Q5 High-Fidelity DNA Polymerase (NEB M0491) and the following primers: 5’-atgGCGGCCGCcagcgg-3’ and 5’-aaaCTCGAGtcagtgccgccgcatcttcacc-3’ (enzyme digestion sites in capital letters). The 5’end of the ALKBH5 coding regions naturally contains a Not I digestion site while the reverse primer introduces a Xho I site right after the stop codon. This PCR product was subsequently digested with Not I and Xho I, and ligated into the matching digestion sites of the lentiviral vector pLEX-FLAG, which encodes double FLAG tags 5’ of the Not I site [20].

Similarly, YTHDC1 was cloned out of cDNA by PCR using the primers: 5’-aaaGCGGCCGCtgcggctgacagtcggg-3’ and 5’-aaaCTCGAGttatcttctatatcgacctctctcccctc-3’ (enzyme digestion sites in capital letters), digested and ligated into the Not I and Xho I sites of pLEX-FLAG, producing pLEX-FLAG-YTHDC1 which encodes an N’-2xFLAG-tagged YTHDC1. This FLAG-YTHDC1 was subsequently PCR clones with primers 5’-aaaAAGCTTatggattataaggatgatgatgataaagg-3’ and 5’-aaaGAATTCttatcttctatatcgacctctctccc-3’, then digested with Hind III and Eco RI and ligated into pK-Myc to build the CMV promoter driven expression vector pK-FLAG-YTHDC1.

### Generation of ALKBH5 overexpression CEM cells

The lentiviral pLEX-FLAG-ALKBH5 vector was packaged in 293T cells and used to transduce CEM cells. At 3 days post transduction, cells were subject to selection with 2µg/mL puromycin for a week until an accompanying untransduced well of cells was completely dead, then the transduced cells were single cell cloned by limiting dilution. Three clones that strongly expressed a FLAG-ALKBH5 protein band slightly larger then endogenous ALKBH5 were used for downstream assays.

### YTHDF2 knockout and over-expression cells

CEM-SS cells with YTHDF2 knocked out by CRISPR/Cas9, overexpressing cells from transduction with pLEX-FLAG-YTHDF2, and the corresponding pLEX-FLAG-GFP transduced control cells, were previously characterized [20].

### Validation of m^6^A depletion in ALKBH5 over-expression cells

4 million each of WT and +ALKBH5 CEM cells were infected with NL4-3 virus, harvested 3 days post infection (dpi), and total RNA extracted with 1ml of Trizol. 150µg of total RNA from each sample were poly(A) purified twice using the Poly(A)Purist MAG Kit (Invitrogen #AM1922). After the first poly(A) purification, the used oligo-dT beads were kept in PBS in tubes marked with the samples they were used on, and subsequently used for the 2^nd^ round of poly(A) purification of the same samples with 30% fresh beads added. All double poly(A) purified RNAs were normalized to a concentration of 75ng/µl, and 200ng of each sample analyzed for m^6^A content using the EpiQuik m6A RNA Methylation Quantification ELISA Kit (EpiGentek # P-9005-48). The results were read using a BMG Clariostar plate reader at the Duke Functional Genomics core facility to read the absorbance at 450nm. To validate the purity of double poly(A) purified RNA, 75ng each of pre- and post-purification RNA were analyzed on a Agilent Bioanalyzer using the RNA 6000 Nano assay, the default analysis software autodetects and quantifies the bands corresponding to the 18S and 28S rRNA.

### HIV-1 replication assays in YTHDF2 overexpressing 293T cells

NLuc virus was packaged in 293T cells in 10 cm plates by transfection of 10µg of pNL4-3-NLuc with PEI (polyethylenimine). For infection of target cells, 293T cells in 6 well plates were co-transfected with 1µg empty vector or pEFtak-FLAG-YTHDF2 [20] and 0.5µg pCMV-CD4 using PEI, media changed 24hrs post, cells passaged 48hrs post, and infected with NL4-3-NLuc 72hrs post transfection, and harvested 24 or 48hrs post infection. Luciferase assays require washing the harvested cells 3 times in PBS prior to lysis in passive lysis buffer (Promega, E1941). NLuc activity was assayed using the Nano Luciferase Assay Kit (Promega, N1120).

### YTHDF reader protein virion packaging assay

5µg pNL4-3-NLuc was co-transfected with 5µg of pEFtak-FLAG-YTHDF1, pEFtak-FLAG-YTHDF2, pEFtak-FLAG-YTHDF3, [20] or pCDNA3-FLAG-agmA3G [25] into 293T cells using PEI, and the cells immediatly treated with 5µM of Indinavir. 3 days post-transfection, producer cells and supernatant media were collected. The supernatant media wwas passed through a 0.45µm pore size filter, and virions purified by centrifugation through a 20% sucrose cushion, followed by banding on a 6-18% Iodixanol gradiant (Sigma D1556) each by 90 mins of ultracentrifugation at 38,000 rpm [19]. Producer cell pellets and purified virion pellets were then lysed and analyzed by Western blot.

### Immunofluorscence microscopy

0.1 million CEM-SS cells were suspended in 300µl PBS and spun onto glass slides using a Thermo Scientific Shandon Cytospin 4 at 1000rpm 3mins. Slides were then immediately fixed in 2% paraformaldehyde in PBS for 10mins, followed by 2 PBS washes. Cells were then permeabilized in 0.3% Triton-X 100 in PBS for 20mins, and washed twice with PBS, and blocked with 1% BSA in PBS for 45mins. Cells were stained with 1/1000 or 1/3000 diluted antibodies targeting YTHDF2 or YTHDC1 (in 1% BSA PBS), over night at 4°C, followed by 3 washes. Cells were then counter stained with 1/1000 diluted Alexafluor 488 (Invitrogen A11034) anti-rabbit, and washed twice with PBS, once each with 70% and 100% EtOH, and air dried briefly. Each cytospin dot was mounted with Vectashield (H1200), pre-mixed with DAPI, and left at −20°C for a few days. Slides were visualized with an EVOS M5000 microscope at 40x magnification.

### YTHDC1 knockdown and overexpression assays

On day 0, 293T cells were seeded at 2 million cells in 8ml DMEM per 10cm plate and 0.4 million cells in 2ml DMEM per 6 well plate well. The next day (day 1), the 10cm plate was transfected with 10µg of pNL4-3 plasmid using PEI. The 6 well plates were used as target cells, with each well transfected with non-targeting (siCtrl) or YTHDC1-targeted siRNAs (siDC1, Origene # SR314128, three YTHDC1 siRNAs premixed at a 1:1:1 ratio): 2.5µl of a 2nmol stock siRNA was diluted in 122.5µl of OPTI-MEM (Gibco), and 7.5µl of Lipofectamine RNAiMax (Invitrogen) was separately diluted in 117.5µl of OPTI-MEM. The pre-diluted siRNA and Lipofectamine solutions were combined and, 5mins later, added drop-wise to the cells in 6 well plates. 6-8hrs post siRNA transfection, the transfection media was removed from 6 well plates, and the cells overlaid with fresh DMEM. The cells in 6 well plates were subsequently co-transfected with 0.5µg pCMV-CD4, 1.5µg pBC-CXCR4, and 1µg of either empty pK vector or pK-FLAG-YTHDC1, using PEI. On day 2, the media of 10cm and 6 well plates were exchanged for fresh DMEM, and the 6 well plates were subject to a second round of siRNA transfection. The transfections were coordinated for a control well transfected with pK vector + siCtrl, a knockdown well of pK vector + siDC1 and an overexpression well of pK-FLAG-YTHDC1 + siCtrl. On day 3, the media was changed again on the 6 well plates to remove the transfection mix. Day 4, the cells in the 6 well plates were trypsinized, counted and seeded either 0.5 million in 0.5ml DMEM per well in 12 well plates or 1 million in 1ml DMEM per well in 6 well plates. The NL4-3 virus-containing supernatant media from the 10cm plates was filtered through a 0.45µm pore size filter to remove any contaminating cells, brought up to 12mls with DMEM, and added to the just seeded infection target cells: 2ml/well of virus for infections in 6 well plates, 1ml/well for 12 well plates. To limit to a single cycle infection, infected cells were treated with 133µM of the reverse transcriptase inhibitor Nevirapine (Sigma SML0097) at 16 hpi, then harvested at 48 hpi for protein or RNA analysis. For each experiment, an aliquot of each siRNA/CD4-transfected infection target cells were seeded in a 12 well plate the day of infection for Western blot validation of YTHDC1 knockdown the next day.

### Subcellular fractionation assays

HIV-1 infected 293T cells, expressing wild type or knocked down levels of YTHDC1, were lysed and divided into nuclear and cytoplasmic fractions as previously described [39].

### Single cycle replication assays in T cell lines

These were performed as previously described [44]. Briefly, NL4-3 virus was packaged in 10cm plates of 293T cells, and the culture media exchanged with fresh RPMI media the day after transfection. 3 days post transfection, the virus-containing supernatant media were harvested and filtered through a 0.45µm pore size filter, and the virus-containing filtrate brought up to a total volume of 12 ml with RPMI. The day before infection, infection target cells of interest were passaged to ensure cells were growing healthily. Infection target cells were counted and seeded at 1 million cells per 12 well plate in 0.5ml fresh RPMI, and subsequently overlayed with 1ml of the virus-containing filtrate. To limit to a single cycle infection, infected cells were treated with 133µM of the reverse transcriptase inhibitor Nevirapine (Sigma SML0097) at 16 hpi, then harvested at 48 hpi. Cells were washed once with PBS prior to lysis with Trizol for RNA extraction or Laemmli buffer for Western blot analysis.

### Western blots

Harvested and PBS washed cells were lysed in Lamalli buffer and kept at −20°C until ready for analysis. Prior to electrophoresis, cell lysates were sonicated till homogenous, denatured at 95°C 15mins, separated on 4-20% or 8-16% gradient Tris-Glycine SDS polyacrylamide gels (Invitrogen), then transferred to a nitrocellulose membrane. Membranes were blocked in 5% milk in PBS + 0.1% Tween (PBST+milk) and stained with primary and secondary antibodies diluted 1:5000 in PBST+milk for 1hr to over night. Signals were visualized by chemiluminescence with 2.5mM Luminol, 0.4mM p-Coumaric acid, 0.1M Tris-HCl pH8.5 in dH_2_O, supplemented with 0.0192% hydrogen peroxide added prior to imaging.

### RNA quantification by qRT-PCR

Done as previously described [44]. 1µg per sample of Trizol extracted RNA were treated with 0.15µl/reaction of DNase I (NEB) and 1µl 10x DNase buffer in a 10µl reaction for 30mins at 37 °C, followed by adding 5µl of 15mM EDTA and inactivating at 75 °C 10mins. 5µl of DNase treated RNA was then used for reverse transcription (RT) with Super Script III (invitrogen) and random hexamer primers following manufacturer instructions in a 20µl reaction at 50°C 1hr followed by 15mins inactivation at 70°C. Each 20µl RT reaction was diluted with dH_2_O to a final 120µl. qPCR was performed with 6µl diluted RT product, 8µl 2x Power SYBR Green Master Mix (ABI), and 1µl each of 1mM forward and reverse qPCR primer in a 16µl reaction. qPCR readouts were normalized to GAPDH levels using the delta-delta Ct (ΔΔCt) method. All PCR primers used are listed in Supplemental Table 1.

### RNA decay assay

RNA stability were measured using the transcription stop method as previously described [44]. Single cycle infected cells were treated at 2 days post infection with 5µg/ml Actinomycin D (Sigma A9415), and harvested 0, 2.5, 5, and 7.5 hrs later. Viral RNA levels at each time point were then assayed by qRT-PCR, calculated as fold change over time 0. Statistical analysis of the rate of RNA decay was done on the log_2_ transformation of each fold change value, using the R suite in the Rstudio interface, comparing slopes of linear regression lines by analysis of covariance (ANCOVA) with the aov function using the following code:

> DATATABLE <-read.csv(“datafile.csv”, header = TRUE)

> modv <-aov(val∼type, data=DATATABLE)

> summary(modv)

### YTHDC1 PAR-CLIP

PAR-CLIP was performed as before with minor modifications [20, 30]. For each immunoprecipitation, 7x 15cm plates of 293T cells were seeded at 5 million cells/plate. The next day, 5 plates were transfected with 19µg/plate of pNL4-3 for virus production, the other 2 plates with (per plate) 2.5µg of pCMV-CD4, 10µg of pBC-CXCR4 and 7.5µg of plasmids expressing either FLAG-GFP or FLAG-YTHDC1, all transfections with PEI. The next day, the virus production plate media was changed, the CD4/CXCR4/FLAG-tagged gene of interest transfected plates were split 1 in 5 to a total of 10x 15cm plates of infection target cells. 3 days post transfection, the supernatant of virus production cells were spun 3000rpm 10mins and filtered through 0.45µm pore filters to remove floating cell, diluted 2x with fresh DMEM, and used to infect the infection target plates, resulting in 15ml media/virus mix per plate. 48hpi, 8ml of supernatant were removed from each plate, and replaced with 10ml of fresh media supplemented with 4-thiouridine (4SU) to a final concentration of 100µM. The next day, 16hrs post 4SU pulse, the supernatents were replaced with 3ml/plate of ice cold PBS, and subject to 2500 x100µJ/cm2 of 365nm UV irradiation twice. Cells were then washed twice with PBS, scrapped off plates, and frozen at −80°C. PAR-CLIP was then performed using a FLAG antibody (M2, Sigma) as described in a published protocol [45]. RIPA buffer (NaCl 150 mM, Nonidet P-40 1%, Sodium deoxycholate 0.5%, SDS 0.1%, Tris-HCl pH 7.4 25 mM) was used in place of PAR-CLIP lysis buffer to ensure sufficient release of nuclear proteins. After elution, the resulting RNA was passed through Biorad Micro Bio-Spin Columns with Bio-Gel P-30 to remove excess salts, then subject to Illumina sequencing library preparation using the Illumina TruSeq Small RNA Library Prep Kit. Multiplex barcode indexes used were RPI21, 22 and 23 for samples GFP control, DC1 repeat 1 and DC1 repeat 2, respectively.

Sequencing was done on a HiSeq 2000 in 50bp single read mode at the Duke Center for Genomic and Computational Biology core facility. Data analysis was done as previously described [44]. Reads <15nt or with a fastq quality score < Q33, along with sequencing adaptors were removed using the FASTX-toolkit v0.014 [46]. Past-filter reads were then aligned to the human genome (hg19) using Bowtie, and the human non-aligning reads then aligned to the HIV-1 NL4-3 sequence with a single copy LTR (U5 on the 5’ end, and U3-R on the 3’ end), thus 551-9626 nt of GenBank AF324493.2.), allowing 1 mismatch. An in-house Perl script was used to only retain reads containing the T>C conversions resulting from UV-crosslinked 4SU. Peak calling was done using PARalyzer v1.1 [47] (with the parameters: BANDWIDTH=3 CONVERSION=T>C MINIMUM_READ_COUNT_PER_GROUP=10 MINIMUM_READ_COUNT_PER_CLUSTER=1 MINIMUM_READ_COUNT_FOR_KDE=5 MINIMUM_CLUSTER_SIZE=15 MINIMUM_CONVERSION_LOCATIONS_FOR_CLUSTER=2 MINIMUM_CONVERSION_COUNT_FOR_CLUSTER=1 MINIMUM_READ_COUNT_FOR_CLUSTER_INCLUSION=1 MINIMUM_READ_LENGTH=10 MAXIMUM_NUMBER_OF_NON_CONVERSION_MISMATCHES=1 EXTEND_BY_READ). After file format conversions using SAMtools [48], all data was visualized using Integrative Genomics Viewer (IGV) [49].

### Primer-ID RNA-seq splicing assay

YTHDC1 knockdown or overexpressing 293T cells treated and infected as mentioned above were performed in 6 well plates, 2 wells per sample were collected and the RNA extracted with Trizol (Invitrogen). The samples were subsequently amplified and sequenced as previously described [31].

### PA-m^6^A-seq

Done as previously [20, 39]. Briefly, 270 million each of wildtype and +ALKBH5 CEM cells were used, each infected with HIV-1 produced from ten 15cm plates of 293T cells and pulsed with 100µM of 4SU one day prior to harvest. Total RNA extracted with Trizol and poly(A) purified using the Poly(A)Purist MAG Kit (Invitrogen #AM1922). After m^6^A immunoprecipitation, Illumina sequencing libraries were prepared using the NEBNext small RNA kit (E7300S), using multiplex barcode indexes RPI5 and 6 for control and +ALKBH5 samples, and sequenced on an Illumina Novaseq 6000 sequencer. Data analysis done as mentioned above for PAR-CLIP.

### Quantification and statistical analysis

All statistical details are listed in the figure legends. All averaged data include error bars that denote standard deviation. All statistical analysis done by Student’s T-test comparing subjects to control, with the exception of RNA decay studies, where the slopes of regression lines were compared by ANCOVA.

### Data availability

Deep sequencing data for the YTHDC1 PAR-CLIP have been deposited at the NCBI GEO database under accession number GSE165473, while the splicing analysis data is under accession number GSE166237.

## Acknowledgements

The following reagents were obtained through the NIH AIDS Reagent Program, Division of AIDS, NIAID, NIH: HIV-1 p24 Gag Monoclonal (#24-3) from Michael Malim and HIV-1 NL4-3 Infectious Molecular Clone (pNL4-3, #114) Malcolm Martin. We thank Ananda Ayyappan Jaguva Vasudevan for cloning assistance, as well as the labs of Christopher Holley, Nicholas Heaton, Dennis Ko and Qi-Jing Li for allowing us to use their equipment.

## Supplemental Material

**Fig. S1.**
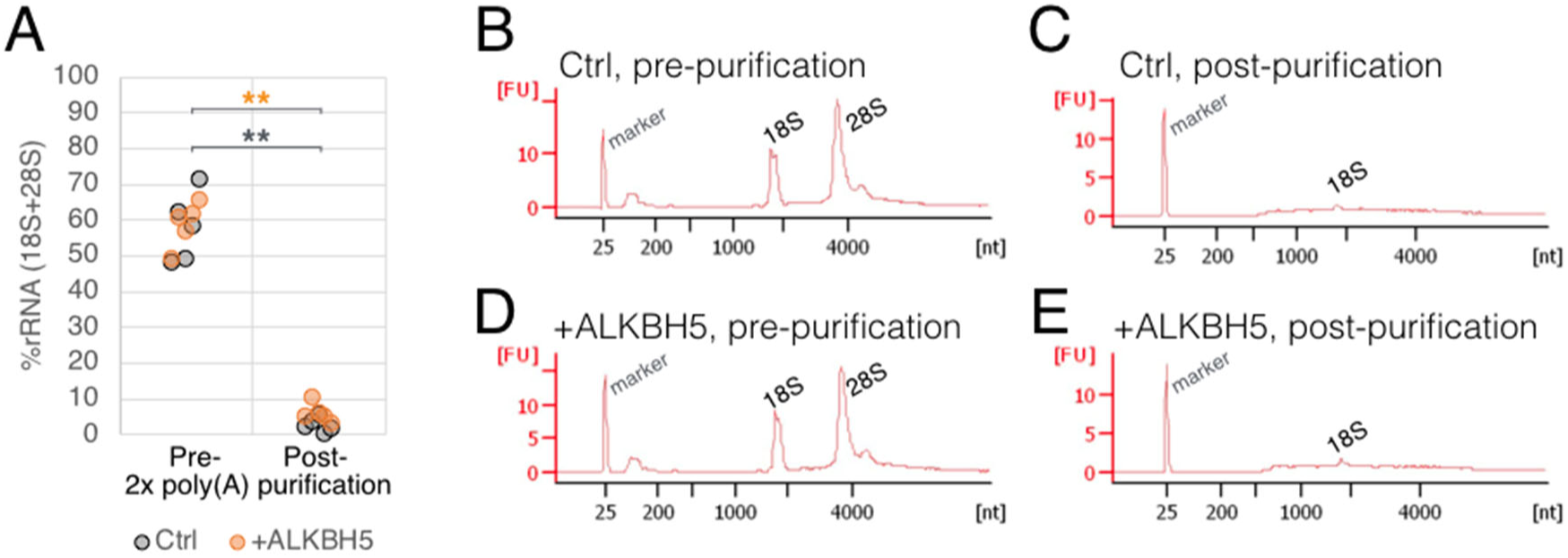
Validation of poly(A) purification for m^6^A ELISA quantification. 75ng each of pre-purification and post-purification RNA from WT CEM T cells (Ctrl) and ALKBH5-overexpressing CEM T cells (+ALKBH5) were analyzed on an Agilent Bioanalyzer electrophoresis system to validate the efficiency of rRNA removal using oligo-dT. (A) Percent rRNA content in each RNA samples from 5 independently collected sets of RNA. (B-E) Representive quantification curves of a sample set used to graph panel A. Peaks corresponding to the expected sizes of 18S and 28S rRNA as labeled.

**Fig. S2.**
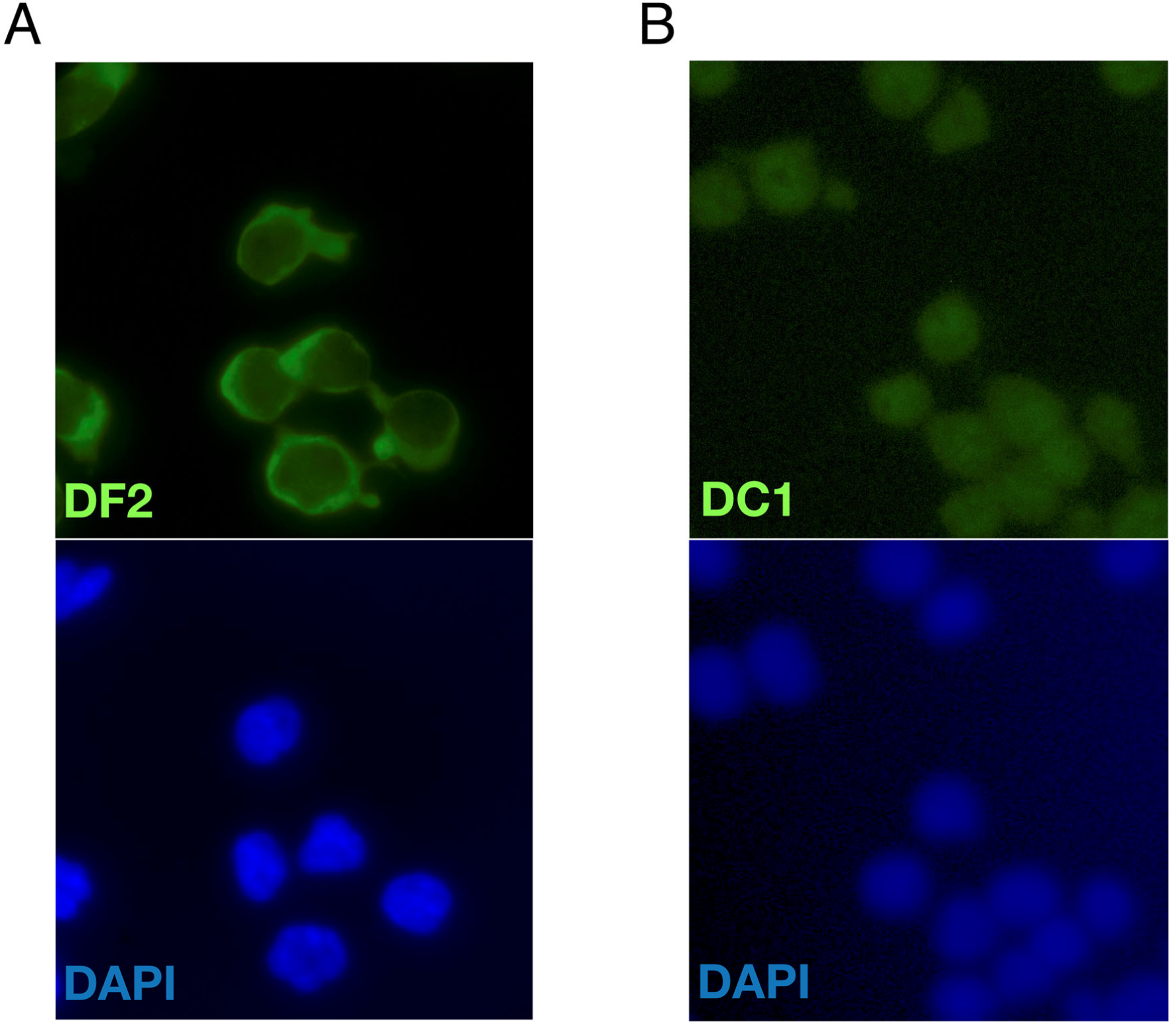
Intracellular localization of m6A readers YTHDF2 and YTHDC1 in CD4^+^ T cell lines. CEM-SS T cells were stained with antibodies targeting endogenous YTHDF2 (A) or YTHDC1 (B), co-stained with the nuclear stain DAPI, then visualized by fluorescence microscopy.

